# RNA BaseCode: Long-Read RNA Sequences with Short-Read Accuracy

**DOI:** 10.64898/2025.12.09.693136

**Authors:** Gert-Jan Hendriks, Anton J.M. Larsson, Paloma Ruiz de Castroviejo Teba, Michael Hagemann-Jensen, Christoph Ziegenhain, Rickard Sandberg

## Abstract

RNA sequencing has revolutionized the life sciences by enabling comprehensive profiling of gene expression across tissues and individual cells. Most mammalian genes produce multiple isoforms with distinct functional consequences, yet their accurate detection and quantification remain challenging. Here we present RNA BaseCode, a synthetic long-read RNA sequencing method that combines the high accuracy and quantitative precision of short-read sequencing with internal random barcodes to reconstruct full-length RNA sequences. We applied RNA BaseCode to purified RNA to benchmark its performance against short-read and long-read RNA sequencing methods. RNA BaseCode demonstrated superior basecalling accuracy, higher quantitative precision, and an improved ability to reconstruct full-length transcripts compared to long-read sequencing. This method enables comprehensive characterization of RNA isoform expression at scale, providing a powerful and broadly applicable tool for investigating isoform regulation across biological contexts as well as the roles of these isoforms in health and disease.

## Introduction

Short-read RNA sequencing remains the gold standard for high-throughput gene expression profiling, offering high accuracy, scalability, and broad accessibility. Application of this technology across human tissues, has revealed that most human genes generate multiple transcript isoforms^1,2.^ Many of these isoforms are functionally distinct^3^ and erroneous splicing of RNA is widely reported to be associated with disease^4^. However, short-read sequencing relies on fragmentation of input RNA (or cDNA) during library preparation. This complicates the quantification of full-length isoforms and instead restricts interpretation to estimation of local splice sites (e.g., miso^5^) or probabilistic inference of transcript structures (e.g., Kallisto^6^ and RSEM^7^). In contrast, long-read technologies can sequence whole cDNA, and in some cases even RNA, enabling direct enumeration of RNA isoform sequences, although at the expense of lower accuracy, quantitative precision, availability and throughput^8–10^.

Molecule identifying barcodes, such as UMIs, are typically located at either end of the cDNA, corresponding to either the 3’ or the 5’ end of the RNA molecule. We realized that inserting molecule-identifying barcodes, consisting of unique error patterns, throughout the first-strand cDNA would allow us to identify short reads sharing the same patterns as originating from the same RNA molecule.

Here, we report RNA BaseCode, a method that uses base analogues with promiscuous base-pairing to introduce molecule-specific base conversions during reverse transcription. (Figure 1a) Thus, each reverse-transcribed cDNA receives a molecule-specific error-code that becomes superimposed onto its original sequence. After full-length cDNA amplification, fragmentation, short-read sequencing, and alignment to the reference genome, the error-code facilitates the reconstruction of original full-length RNA transcripts in a quantitative and highly parallel manner. As each cDNA is reconstructed from several overlapping short-read sequences, RNA BaseCode achieves high base-calling accuracy, precise exon junctions, and eliminates most PCR-induced errors as well as virtually all sequencing errors in the reconstructed RNA sequences. RNA BaseCode combines the benefits of short-read sequencing with superior full-length RNA isoform reconstruction and quantification compared to conventional long-read sequencing approaches

**Figure 1.**
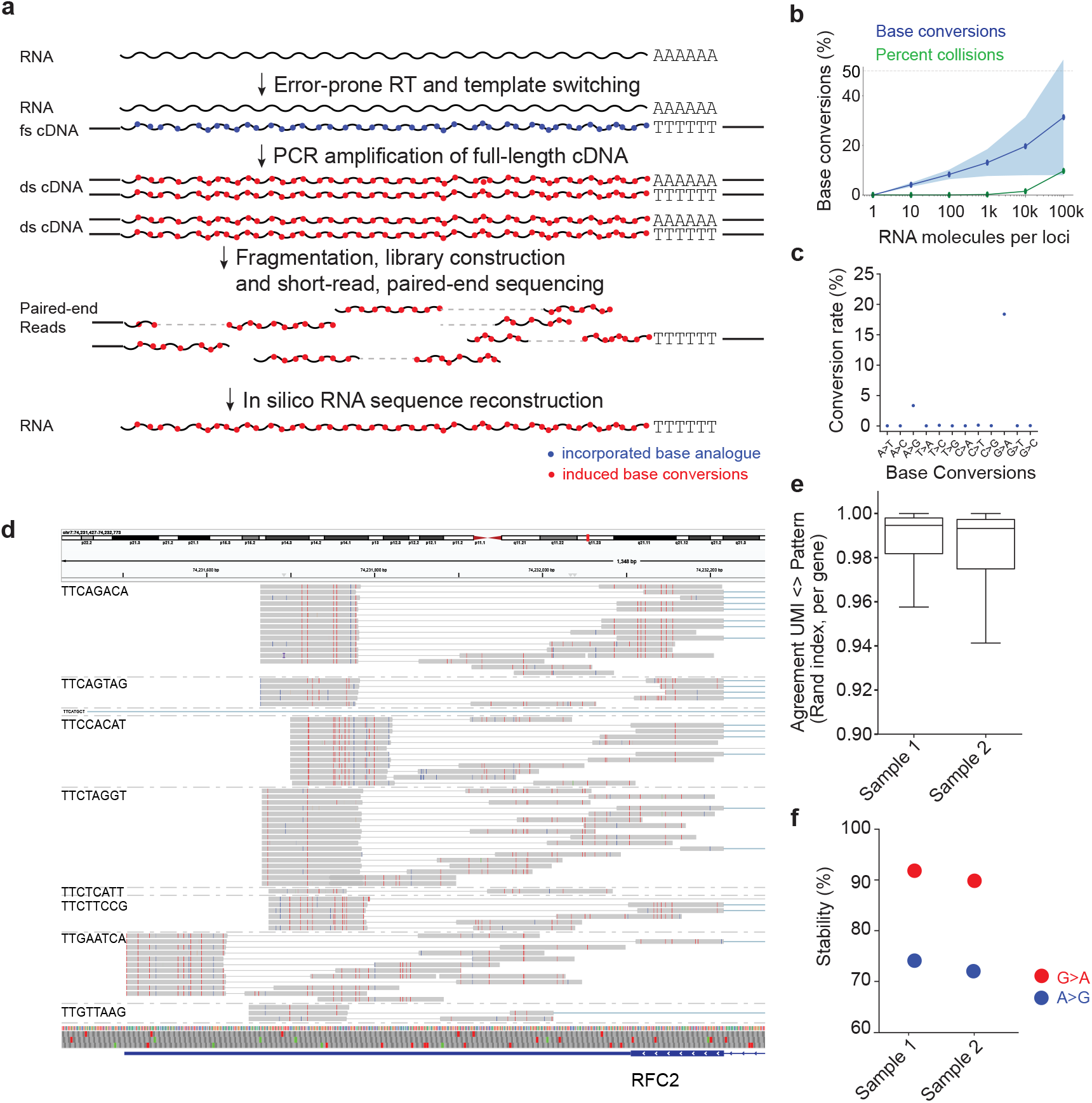
RNA BaseCode synthetic long-read RNA sequencing concept. **(a)** Illustrating the RNA BaseCode synthetic long-read RNA sequencing technology. RNA is subject to reverse transcription in the presence of a base analogue that drives base conversions. Each transcribed first-strand cDNA obtains a unique pattern of incorporated base analogues that, in appropriate subsequent reaction conditions, lock in a pattern of base conversions in the corresponding cDNA sequence. Full-length cDNA is subjected to fragmentation-based indexing followed by short-read sequencing. Paired-end short reads are computationally reconstructed based on the unique error-code present in each original cDNA molecule. **(b)** Simulation showing the base conversion percentages required to distinguish increasing numbers of RNA molecules per gene loci. Blue data points show the mean base conversion percentage with shaded region showing the 95% confidence interval. Green data points show the percentage of collisions (mean and confidence interval), i.e., identical error-patterns in sequence overlaps that originated from different original RNA molecules that could lead to erroneous reconstructions. **(c)** Dotplot showing the percentage of induced G-to-A base conversions. **(d)** Showing aligned short 3’ reads from the RFC2 gene in RNA extracted from the HEK293FT cell line, separated by their unique molecular identifier (the UMI sequence is shown to the left). Each RNA has a dense and unique error pattern. **(e)** Boxplot showing agreement between reads grouped by UMI sequence and pattern of base conversions. **(f)** Estimated stability of G-to-A and A-to-G base conversions in two representative RNA BaseCode samples.

## Results

We first used simulations to determine the number of base conversions required to distinguish tens of thousands of RNA molecules per gene locus using a base analogue that introduces one specific base conversion (and assuming 100 bp short-read overlaps). The analysis showed that a 10% base conversion rate separates up to 100 distinct RNA molecules per locus, where a 20% rate can distinguish up to 10,000 RNA molecules and with conversions rates around 30% it is feasible to distinguish 100,000 distinct RNA molecules (Figure 1b). These error rates are orders of magnitudes higher than those obtained with reverse transcriptase enzymes, demonstrating the need to develop novel strategies for efficient introduction of errors into cDNA during reverse transcription.

To overcome these limitations, we established the RNA BaseCode platform for synthetic long-read RNA sequencing as a robust and easy-to-use RNA sequencing library preparation method that produces high rates of G>A base conversions (Figure 1c). During cDNA library preparation, RNA is reverse transcribed in the presence of the promiscuous base analogue dPTP.

This base analogue can basepair with both guanine or adenine bases and in appropriate reaction conditions this will result in high rates of G>A base conversions that uniquely barcode each cDNA molecule. Thereafter, enzymatic cleanup and template switching are performed in separate reactions. After first-strand cDNA clean-up, the second strand is synthesized, and the resulting material amplified. From this point, sequencing libraries are prepared using an established fragmentation and ligation approach.

To standardize coverage over the RNA molecules, fragments corresponding to the 3’ end of the RNA molecule are separately amplified from fragments corresponding to the 5’ or internal part of the RNA molecule. To validate that the base conversion patterns reflect and quantify individual RNA molecules, we introduced a unique molecular identifier (UMI) into the oligo-dT and compared the UMI sequence to the observed patterns. After aligning the sequenced short-read data against the reference genome, we found high concordance between the UMI sequence and the observed patterns (Figures 1d and 1e). To confirm that the pattern is stable in the amplified cDNA molecules originating from the same first-strand cDNA molecule, we then confirmed their per-conversion stability. The introduced patterns were highly stable (>90%) for G>A base conversions and less stable (75%) for A>G base conversions (Figure 1f), presumably related to the relative kinetics of incorporation of adenine and guanine opposite a P template^11^. We conclude that error-prone reverse transcription, using base analogues with promiscuous base-pairing, can induce abundant base conversions in cDNA uniquely identifying the RNA molecule of origin that can be used for sequence reconstruction.

Next, we generated data for four samples of total RNA from HEK293FT cells (5 ng each) that were sequenced to a depth of approximately 50 million 150-bp paired-end reads per sample. The data was then processed using a software pipeline specifically developed to address the unique properties of RNA BaseCode (Figure S1, methods). Aligning the shortread data to the human reference genome demonstrated consistent, specific, and high-density base conversion patterns with a mean of 22% of Gs converted into A (Figure 2a). Since base conversion errors for genes located on the negative DNA strand are detected as C-to-T conversions (Figure 2a), the identity of the induced base conversions is sufficient to determine strandedness for all individual paired-end reads and the reconstructed RNA molecules (Figure 2b). Furthermore, we observed that the introduced error patterns are not affected by local sequence context, confirming that the simulations (shown in Figure 1) are an accurate estimation of our power to distinguish thousands of distinct RNA molecules originating from the same gene based on their unique error patterns (Figure S2).

**Figure 2.**
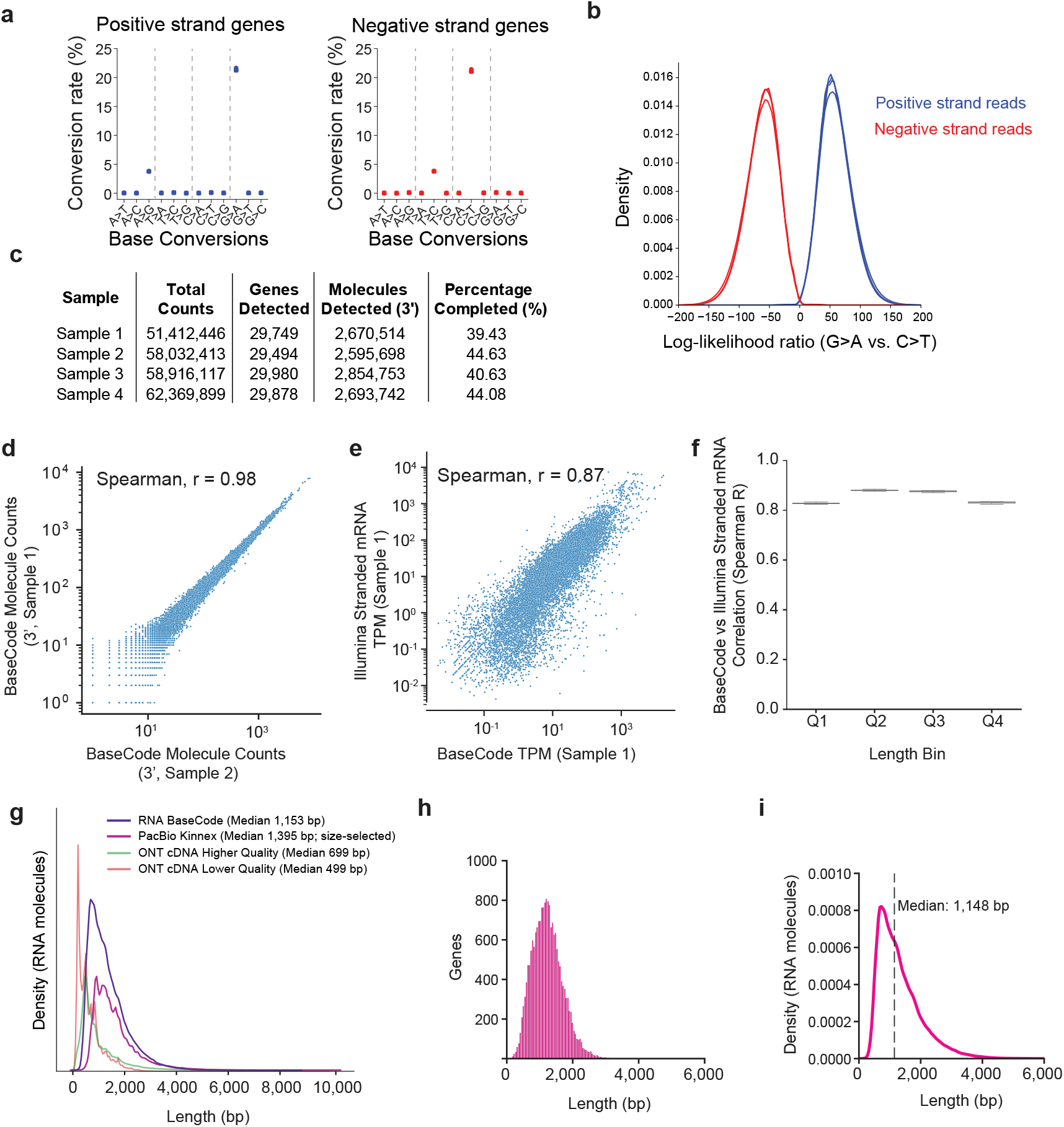
Benchmarking BaseCode on HEK293FT RNA. Showing the specific induction of G-to-A (and to a smaller degree A-to-G) base conversions in RNA BaseCode samples, separating for reads originating from genes on the positive and negative strand. **(b)** Log-likelihood distributions of paired-end reads to have originated from the positive and negative strand, colored according to originating from a gene on the positive and negative strand. **(c)** Table summarizing the counts of reads, detected genes, detected RNA molecules and completely reconstructed RNA molecules per sample. **(d)** Scatter plot comparing RNA molecule counts from two representative RNA BaseCode experiments across 18,142 protein-coding genes. **(e)** Scatter plot comparing transcript per million (TPM) expression values estimated by RNA BaseCode (x-axis) and Illumina Stranded mRNA (y-axis) across 18,142 protein-coding genes. **(f)** Pairwise spearman correlation between RNA BaseCode and Illumina Stranded mRNA samples within protein-coding gene groups binned by transcript length (Q1: 0-1.25 kb, Q2: 1.25-2 kb, Q3: 2-3.2 kb, Q4: 3.2 kb+, n = 4,535 or 4,536 genes per bin. **(g)** Length distribution of molecules sequenced on either long-read platforms or reconstructed using RNA BaseCode, colored by technology and showing their respective median length. **(h-i)** Histogram of the mean length of completed reconstructed molecules across 29,749 genes for Sample 1 **(h)** and the length distribution of the individual completed reconstructed molecules **(i)**.

We reconstructed the paired-end short-read data into synthetic long reads (methods). Between 29,494 and 29,980 genes (Ensembl gene annotations) were detected in all samples, corresponding to more than 2.5 million captured molecules per sample based on the number of unique error patterns observed in 3’ reads (See table in Figure 2c). Of these molecules, approximately 40% were reconstructed from the 3’ end to the 5’ end, resulting in completely reconstructed molecules (see table in Figure 2c). An additional 25% of 3’-anchored molecules were reconstructed by the inclusion of internal reads but did not contain 5’ reads, indicating that these are partially reconstructed molecules. Importantly, the RNA BaseCode protocol was highly reproducible between the HEK293FT samples, with all sequenced samples having highly concordant 3’-based molecule counts across genes (Figure 2d, Figure S3, spearman r = 0.98, protein-coding genes).

To understand the extent to which RNA BaseCode quantifies gene expression consistent with established short-read RNA-sequencing methods, we prepared and sequenced four Illumina Stranded mRNA libraries from the same HEK293FT RNA (methods). These libraries were analyzed using software as similar as possible to the RNA BaseCode pipeline to reduce discrepancies due to the computational methods used (methods). The Illumina Stranded mRNA libraries were also highly reproducible in an intra-method comparison (Figure S4, Spearman r = 0.97-0.98, protein coding genes). Importantly, the gene expression estimates between Illumina Stranded mRNA and RNA BaseCode were in quantitative agreement in all cross-method comparisons (Figure 2e, Figure S5, Spearman r = 0.87, protein coding genes). The agreement was consistent across genes with different transcript lengths (Figure 2f).

We next investigated the performance of the synthetic long-read reconstructions achieved with RNA BaseCode. Comparing these results to cDNA typically obtained on long-read sequencing platforms, Oxford Nanopore Technologies (ONT) and Pacific Biosystems, revealed marked differences in length distributions (Figure 2g). In agreement with several reports^8–10^, the mapping lengths of ONT cDNA data were considerably shorter than PacBio Kinnex data. RNAs reconstructed with RNA BaseCode were significantly longer than those from ONT, since median cDNA lengths in ONT data were in the range of 500 to 700 base pairs^9^. Compared to RNA BaseCode, Pacbio Kinnex data had longer median lengths (1.5-1.7 kbp)^8^, however, the increased length is the result of substantial size-selection, resulting in libraries that are heavily skewed towards longer isoforms impairing the accurate quantification of RNAs^8^. The reconstructed lengths obtained for RNA BaseCode varied depending on the gene, reflecting the expected variation in the transcriptome (Figure 2h). For completely reconstructed RNA molecules, their median length was 1,148 base pairs (bp) (Figure 2i), with partially reconstructed molecules (those lacking a 5’ read) having a median length of 735 bp.

Importantly, the RNA BaseCode reconstruction performance was consistent across samples (Figure S6). We conclude that synthetic long-read RNA-sequencing using RNA BaseCode has much improved lengths when compared to cDNA sequencing with ONT, while simultaneously avoiding the length bias introduced during PacBio Kinnex library preparations. RNA BaseCode, however, is quantitative across the whole length span of the transcriptome (Figure 2f).

Finally, we investigated the amounts of artifactual small insertions and deletions (indels) present in long-read sequencing data (ONT) and in reconstructed RNA BaseCode data. Whereas ONT cDNA has many indels per sequence (Figure S7), complicating alignments towards exon-exon junctions, reconstructed RNA BaseCode sequences are devoid of indels and have exact exon-exon junctions. RNA BaseCode reconstructed sequences can occasionally lack internal coverage (exemplified in Figure S7) although typically without any negative impact on isoform assignment of the otherwise fully reconstructed sequence. The lack of indels throughout transcript structures strengthens analyses of small exons and regulated splice site usage that is common among human genes^1^.

## Discussion

Synthetic long-read technologies have been developed for DNA sequencing, where the task is to separate two haplotype sequences. In contrast, synthetic long-read RNA sequencing presents a fundamentally different challenge: hundreds to thousands of distinct RNA molecules can arise from a single gene and each of these needs to be reconstructed independently and without the ability to exploit haplotype-specific genetic variation to distinguish between molecules. Instead, each RNA-derived cDNA must be uniquely barcoded to enable accurate reassembly from short-read data. Synthetic long-read RNA-sequencing strategies have been proposed previously by either separation of cDNA into specific reactions^12^ or by bisulfite treatment of cDNA^13^, prior to fragmentation, barcoding and short-read sequencing. These solutions are not scalable and do not work well since they dramatically reduce throughput or cause cDNA fragmentation respectively.

Several recent studies have discussed strengths and limitations with conventional long-read RNA sequencing approaches, regarding read lengths, biases, accuracy and sensitivity^8–10^. Specifically, ONT data is shorter than expected from typical human transcriptomes, has relatively lower base accuracy and contains numerous indels^14^. PacBio data on the other hand can vary in length and is biased towards long RNA molecules, likely as a result of substantial size-selection^15^. This bias creates substantial challenges when comparing abundances of isoforms of different length for any given gene. To overcome these challenges, long-read RNA sequencing is supplemented with standard short-read RNA sequencing. In addition to being more costly and time-consuming, the combination of two distinct methods also introduces more analytical challenges such as data integration and consumes more of the RNA sample. With RNA BaseCode we obtained both high quantitative accuracy and rich qualitative isoform information while using 60-fold less RNA input than PacBio and 100-fold less than ONT.

The synthetic long-read approach described here leverages consensus calling across short reads originating from the same cDNA molecule. As such, sequencing errors and the majority of PCR errors are removed, with the exception of the molecule-specific pattern of errors that is introduced. This approach we detail here is generally compatible with short-read sequencing platforms and validated on Illumina NovaSeqX, MGI G400 and MGI G99 sequencers. These platforms provide exceptional throughput, data quality, operational robustness, and ease-of-use, while being far more accessible to most labs than long-read sequencers.

RNA BaseCode is a platform chemistry that relies on error-prone reverse transcription as a means of internally barcoding cDNA molecules. Importantly, the procedure is in principle compatible with reverse transcription performed in droplets or even in situ reverse transcription for spatial transcriptomics^16,17^, where this strategy would directly enable RNA isoform detection.

## Conclusion

This work introduces RNA BaseCode as a new synthetic long-read RNA sequencing solution that combines the strengths of short-read sequencing, accuracy and throughput, with the ability to resolve full-length RNA isoform. It produces quantitative, high-quality synthetic long reads from as little as 5 ng of total RNA from cell lines, primary tissues, and even RNA purified from cellular compartments. Given the high data quality, low input requirements, and compatibility with widely available short-read sequencing infrastructure, RNA BaseCode is positioned to accelerate RNA and RNA isoform research and deepen our understanding of RNA regulation in health and disease.

## Data Availability

Generated sequencing data has been uploaded and is available through the European Nucleotide Archive (PRJEB105067).

## Code Availability

The complete RNA BaseCode analysis pipeline has been implemented using Snakemake and can be run using the available docker container (https://hub.docker.com/r/basicgenomics/basecode). Moreover, a Snake-make pipeline to generate all the necessary genome reference and annotation files can be found on github (https://github.com/BasicGenomics/Base-CodeGenerate/tree/main). Finally, a small helper script written in BASH is made available to guide a user to run the docker implementation of the pipeline (https://github.com/BasicGenomics/BaseCodeHelper).

## Acknowledgements

We are grateful for grants 2022-02236 and 2023-02946 from Vinnova, the Swedish Innovation Agency, to Basic Genomics and grant 2017-01062 from the Swedish Research Council (to R.S.). We thank Karolinska Institutet Innovation (technology transfer office) for support and mentorship and the Basic Genomics team for critical input to the manuscript.

## Author contributions

GJH, AJML and RS conceived and developed the synthetic long-read RNA-sequencing strategy. GJH was leading the experimental development of RNA BaseCode. GJH and PRdCT generated experimental data. AJML was leading the computational development of RNA BaseCode computational processing and RNA molecule reconstructions. MHJ contributed to library construction strategies. CZ contributed to sequence processing. GJH, AJML and RS analyzed data and wrote the manuscript.

## Competing interest statement

GJH, AJML and RS have filed patent applications and/or are listed as inventors on patent applications relating to RNA BaseCode and co-founded Basic Genomics AB to commercialize the technology. GJH, AJML, and PRdCT are employees of Basic Genomics AB, and GJH, AJML, PRdCT, MHJ, CZ and RS hold equity or options in Basic Genomics AB. BaseCode is a trademark belonging to Basic Genomics AB.

## Materials and Methods

### Culturing and RNA extraction from HEK293FT cells

HEK293FT cells were cultured using standard adherent cell culture conditions. Cells were grown in DMEM (high-glucose) with added 10% FBS, 1% Pen/Strep and sodium pyruvate (1mM). Cells were incubated at 37°C and passaged at ∼90% confluency using TryplE(Gibco). For RNA collection, cells were detached using TryplE, centrifuged at 300x g, washed with PBS, and counted using a Countess 3 system (Thermo). RNA was then isolated using RNeasy Mini Kit (Qiagen) according to the manufacturer’s instructions. The optional DNAse treatment was not performed. The resulting RNA was quantified using a Qubit 4 system with the Qubit RNA high-sensitivity kit. RNA quality was confirmed by capillary gel electrophoresis using the Agilent Bioanalyzer 2100 System and RNA 6000 Nano kit according to the manufacturer’s instructions.

### Preparation of RNA BaseCode sequencing libraries

RNA BaseCode libraries were prepared using Basic Genomics’ RNA Base-Code kit according to themanufacturer’s instructions. In short, reverse transcription was performed in the presence of dPTP. Enzymatic clean-up, and separate template switching reactions were performed, and libraries were cleaned up using the DNA Clean & Concentrator-5 kit (Zymo Research). Second-stranding and amplification were performed, and the resulting amplified cDNA library was then quantified using the Qubit DNA high-sensitivity kit before capillary gel electrophoresis using the Agilent Bioanalyzer 2100 System and the Agilent High Sensitivity DNA Kit. 10 ng of cDNA were used to prepare the final sequencing library. Following the manufacturer’s protocol, fragments corresponding to the body and 5’ part of the RNA molecule are separately amplified from fragments corresponding to the 3’ part of the RNA molecule. These are referred to as PCR A and PCR B respectively. Quantification by Qubit and capillary gel electrophoresis were performed for all resulting libraries. The resulting libraries from PCR A and B from multiple samples were pooled separately and circularized using the MGIEasy Universal Library Conversion Kit (App-A). Pool A and pool B were then pooled to make up 75% and 25% of the final circularized library respectively. This final library was then sequenced, paired-end with 150 cycles for each read and 8 cycles for each index read on the MGI G400 according to manufacturer’s instructions.

### Preprocessing and reconstruction of RNA BaseCode sequencing data

First, the Fastq files were processed to identify the sample and whichpart of the cDNA the read originated from. All index sequences within a hamming distance of 1 were assigned to the identified sample, and all other reads were discarded. The start of read 1 was compared to template switching oligonucleotide (TSO) sequence, TCTTCTCTCCTCCTCC, and each read pair matching this sequence within a hamming distance of 4 was assigned as 5’reads. Reads which were not assigned as 5’-reads were assessed whether the first 24 bases in read 1 contained at least 16 Ts. These reads were assignedas 3’-reads. If the read pair did not pass either check, both read 1 and read 2 were scanned for the TSO sequence using a moving window, and finally both were scanned for the adapter sequence found immediately prior to the T-stretch in 3’-reads, AGATGTGTATAAGAGACAG, to identify additional 5’ and 3’ reads respectively. All read pairs which did not pass any of these criteria were assigned as internal reads. All sequences associated with introduced primers (e.g., the TSO sequence, T stretch, adapter sequence) were trimmed.

The reads were then trimmed additionally using cutadapt(v 4.6) using the command cutadapt –j {threads} –-json{log.summary} –n 2 –m 25 –q 10 –a CTGTCTCTTATACACATCT –a AGATCGGAAGAGCACAC-GTCTGAACTCCAGTCA –g TTTTTTTTTTTTTTTTTTTT –g GGAGGAG-GAGAGAAGA –g AAAAAAAAAAAAAAAAAAAA –A CTGTCTCTTATA-CACATCT –A AAAAAAAAAAAAAAAAAAAA –A AGATCGGAA-GAGCGTCGTGTAGGGAAAGAGTGT –-too-short-output {output.r1_short} –-too-short-paired-output {output.r2_short} –o {output.r1} –p {output.r2} {input.r1} {input.r2}. The trimmed reads were then mapped to the human reference genome hg38 using HISAT-3N using splice sites extracted from the ENSEMBL gene annotation (last updated 2018-07) with the command hisat-3n –-new-summary –-summary-file {log.summary} –k 5 –max-seeds 8 –-score-min L,0,-0.5 –-base-change G,A –-no-temp-splicesite –p {params.cores_hisat} –-known-splicesite-infile{params.splicesites} –x {params.genomeref} –1 {input.r1} –2 {input.r2} | samtools view –F 256 –b –@ {params.cores_samtools} –o {output}. The reads were then written to separate BAM files based on the strand inferred by HISAT-3N, reads with anin-conclusive inferred strand were written to a third BAM file. Each read was then assigned togenes using feature Counts (v 2.1.1), for each strand separately and to all genes for the inconclusive reads.

Molecules were then reconstructed by comparing each overlapping portion between reads and combined into reconstructed molecules if their adjusted mutual information score is above 0.3 (based on the number of covered As, shared number of mutations, and the number of exclusive mutations). If there are multiple possible matches, the empirical false positive rate is calculated for each match. If there is a match which is at least 10 times less likely to be a false positive compared to all other matches, that read is assigned to that molecule. If there are no matches, the read pair will be assigned a new molecule ID and used for future matches.

The reconstructed reads were then flattened by extracting all sequences and PHRED quality scores belonging to the same reconstructed molecule. The sequence at each position was determined by calculating a new likelihood score based on the combined transformed quality scores for each nucleotide and picking the mostly likely nucleotide. The new likelihood score was then converted back into PHRED quality scores. Reference skip (i.e., splicing) was determined by consensus across reads. Missing coverage was assigned the deletion CIGAR symbol due to limitations in the SAM specification.

### Estimation of base conversion pattern stability

To estimate base conversion rates, all reads were compared to the reference genome hg38. All sequenced bases with a read quality below 15 were not considered. All possible mismatches were counted, together with the number of reference bases for each canonical nucleotide (A,T,C,G). The number of mismatches divided by coverage over the reference base was considered as the base conversion rate. The log-likelihood ratio for strandedness was calculated for each read pair by extracting the number of G>A and C>T mismatches to the reference and the number of covered A and T nucleotides, with the requirement that the read matches to an assigned geneas assigned by feature Counts. The log-likelihood ratio was then calculated by subtracting the sum of the likelihood of observing the number of G>A and C>T given that the read originated from a gene on the positive strand by the sum of the likelihood of observing the number of G>A and C>T given that the read originated from a gene on the negative strand. Consequently, reads originating from genes on the positive strand will have a positive log-likelihood ratioand reads originating from genes on the negative strand will have a negative log-likelihood ratio.

To estimate pattern stability in the datasets containing a UMI, reads were grouped by gene and UMI sequence. Each converted position found within each group was extracted, and all positions where a G>A or A>G conversion was found in more than one read was used to estimate stability of that conversion type. Briefly, the number of conversions and the coverage over each considered position was used to calculate a binomial proportion confidence interval using the Wilson method, with the point estimate being the sum of observed conversions divided by the sum of covered positions across reads.

### Preparation of Illumina Stranded mRNA libraries

Illumina stranded mRNA libraries were prepared according to manufacturer’s instructions. For samples 1 and 2, 25 ng total RNA from HEK293FT cells was used. For samples 3 and 4 125 ng of total RNA was used. The resulting libraries were circularized and sequenced on the MGI G99 platform.

### Preprocessing of Illumina Stranded mRNA libraries and comparison to RNA BaseCode

The Illumina Stranded mRNA sequencing reads were processed using nf-core/rnaseq (https://nf-co.re/rnaseq/3.14.0/) using the same reference genome and gene annotation (gff3) file used for RNA BaseCode, first choosing STAR –> RSEM as the alignment and quantification options. The reads were then additionally mapped using HISAT2 and reads annotated by feature Counts, for comparison with RNA BaseCode.

For the comparison between Illumina Stranded mRNA libraries and RNA BaseCode, the RNA BaseCode reads were quantified by counting each fragment assigned toeach gene by feature Counts and then normalizing by the transcript length for that gene as obtained by RSEM from the Illumina Stranded mRNA libraries to obtain FPKM values for each gene, and then further normalized per sample to obtain TPM. The Illumina Stranded mRNA libraries were processed in an identical manner. The gene type (e.g., protein coding) information used for the comparison was obtained from BioMart.

### Comparisons to ONT and PacBio

ONT data was obtained from a recent benchmarking study^9^, and the PacBio Kinnex data was obtained from PacBio (https://downloads.pacbcloud.com/public/dataset/Kinnex-full-length-RNA/DATA-Revio-UHRR/2-FLNC/flnc-1.bam).

For the ONT data, the samples SGNex_K562_cDNA_replicate1_run3 and SGNex_Hct116_cDNA_replicate3_run3 were used, and the number of covered positions for each read is what is used in the comparison. For the visual comparison between ONT and RNA BaseCode for the gene RFC2, SGNex_K562_cDNA_replicate1_run3 and all regular RNA BaseCode samples were loaded into IGV (v. 2.19.2) and mismatched bases were hidden.

For the Kinnex data, the unmapped FLNC reads were used for the length comparison. This is since the pipeline used for PacBio Kinnex data does not produce individual mapped reads, opting instead to immediately construct consensus-level isoform sequences. The presented lengths can therefore be considered the upper bound of the coverage which correspond to actual captured RNA.

For RNA BaseCode, all reconstructed molecules with at least 1 3’-reads, 1 internal read and 1 5’-read (completed reconstructions) across all regular samples were used for the comparison

