## Supplemental Figures for "RNA BaseCode: Long-Read RNA Sequences with Short-Read Accuracy"

**Supplemental Figures for Hendriks, Larsson & Ruiz de Castroviejo Teba *et. al.***

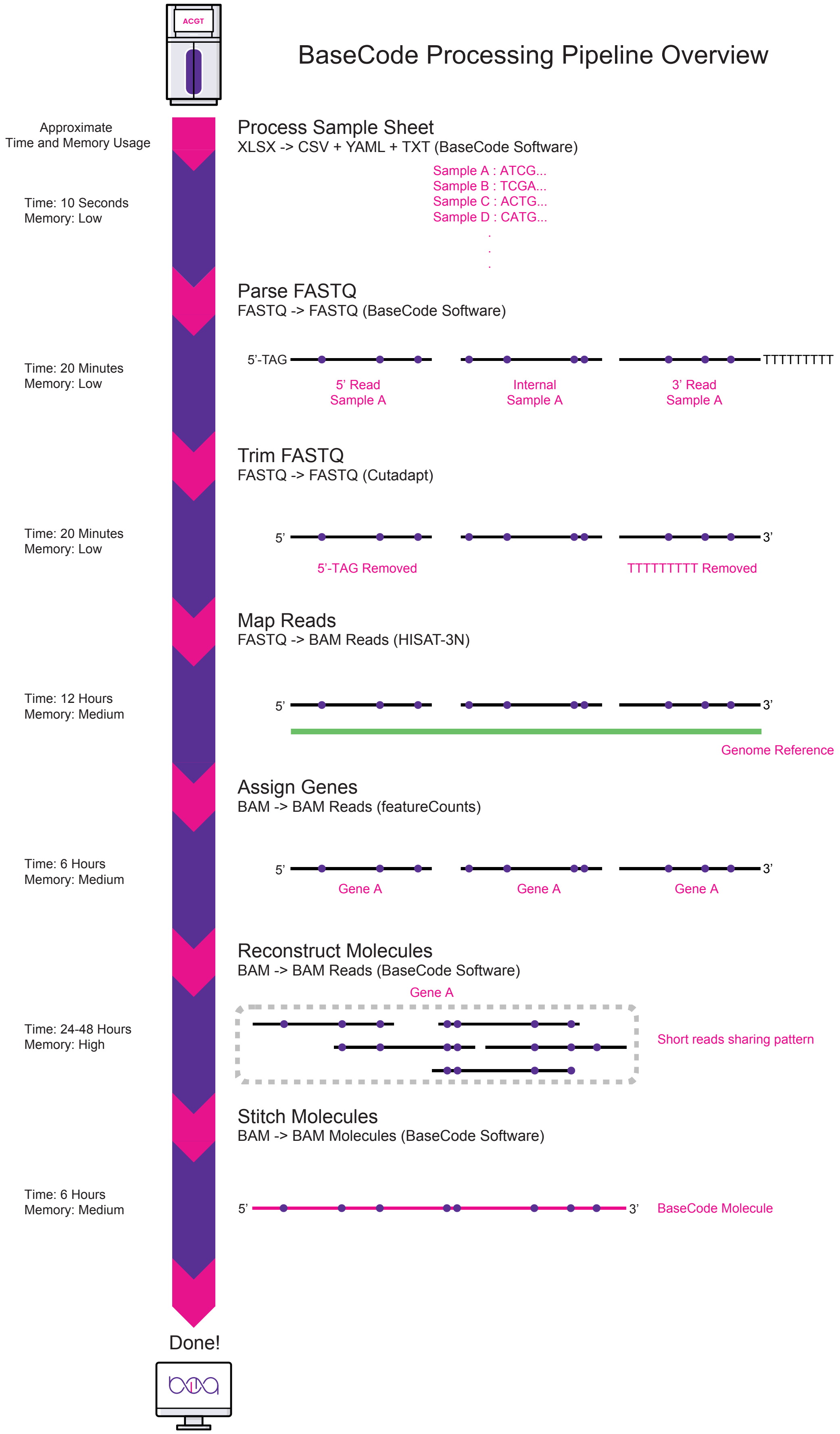


**Figure S1: Overview of the RNA BaseCode data processing workflow. (A)**. An overview of the RNA BaseCode Data processing workflow. Please see the methods and code availability statement for detailed information. Time and resources usage are estimates based on processing datasets for 8 Human RNA samples concurrently.

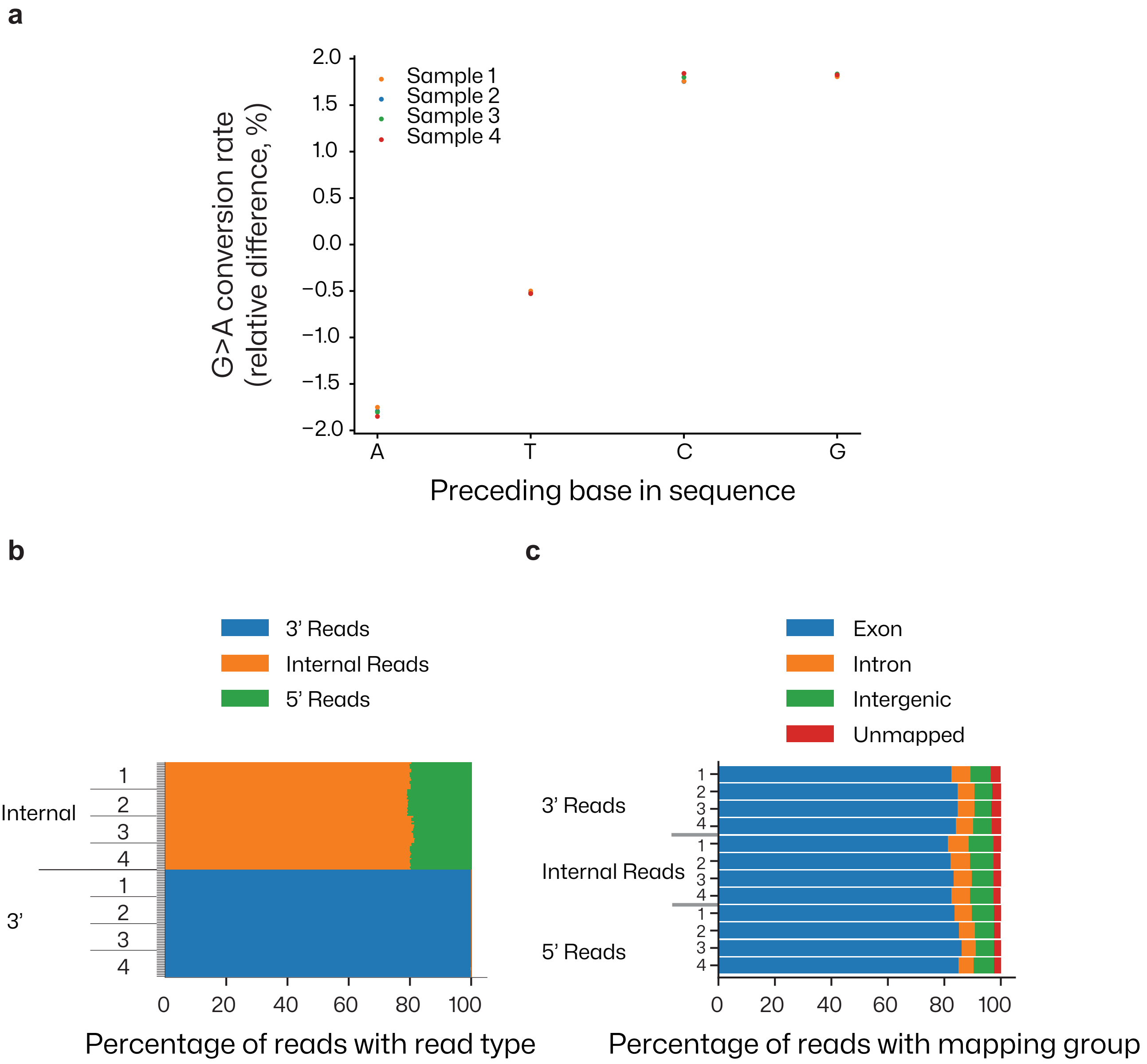


**Figure S2: Quality control metrics for RNA BaseCode data. (a)** Dotplot showing relative local context independence of base conversion rate based on the preceding base. **(b)** Horizontal barplot showing the percentage of reads containing either the 5' or 3' end of original RNA molecule, or internal reads, for each of the the 4 RNA BaseCode experiments. **(c)** Horizontal barplot showing the percentage of reads aligning within exon, intronic, intergenic regions and those that did not align at all (unmapped).

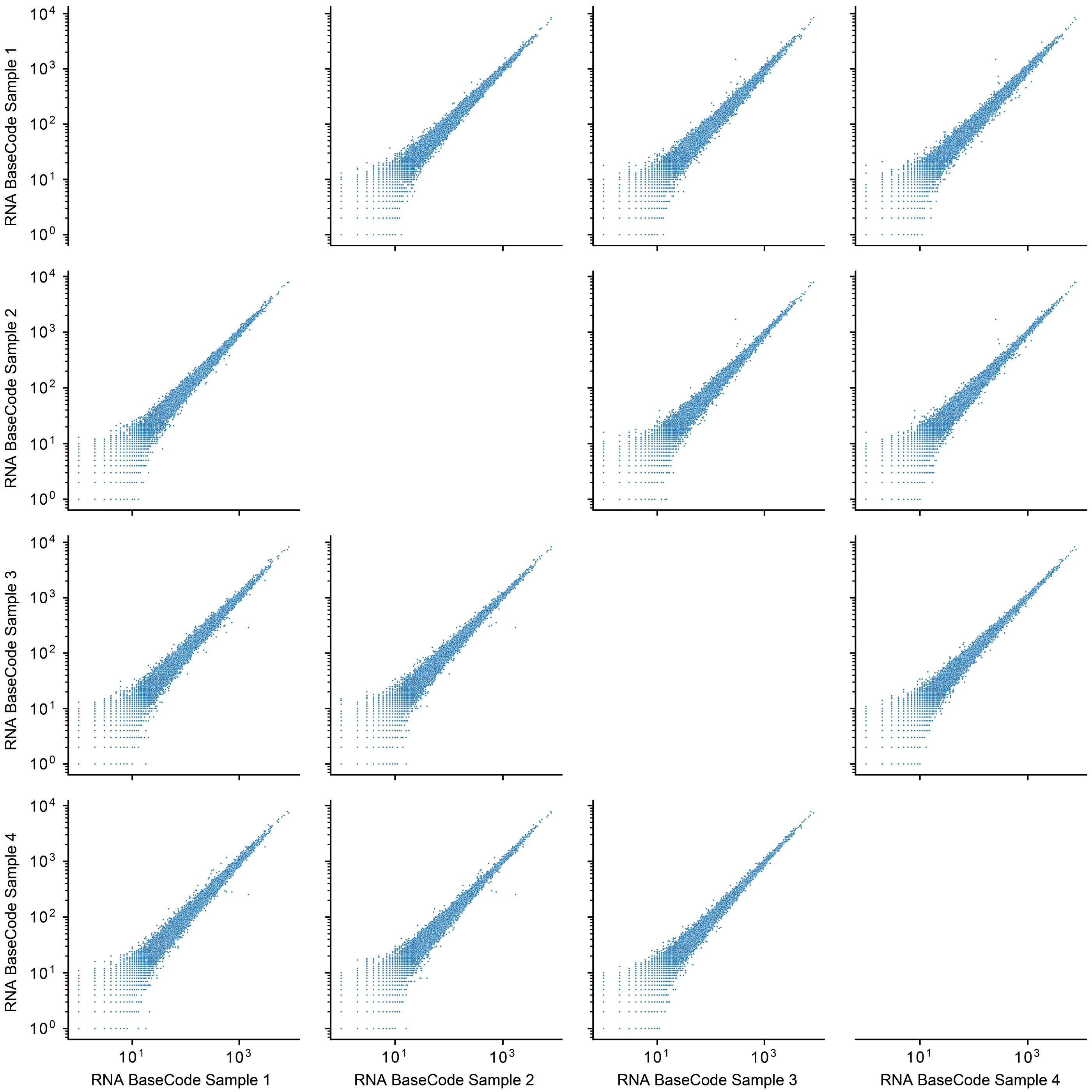


**Figure S3: RNA BaseCode data shows high intra-method correlation.**
Scatter plots of RNA molecule counts from four RNA BaseCode experiments (HEK293FT RNA).

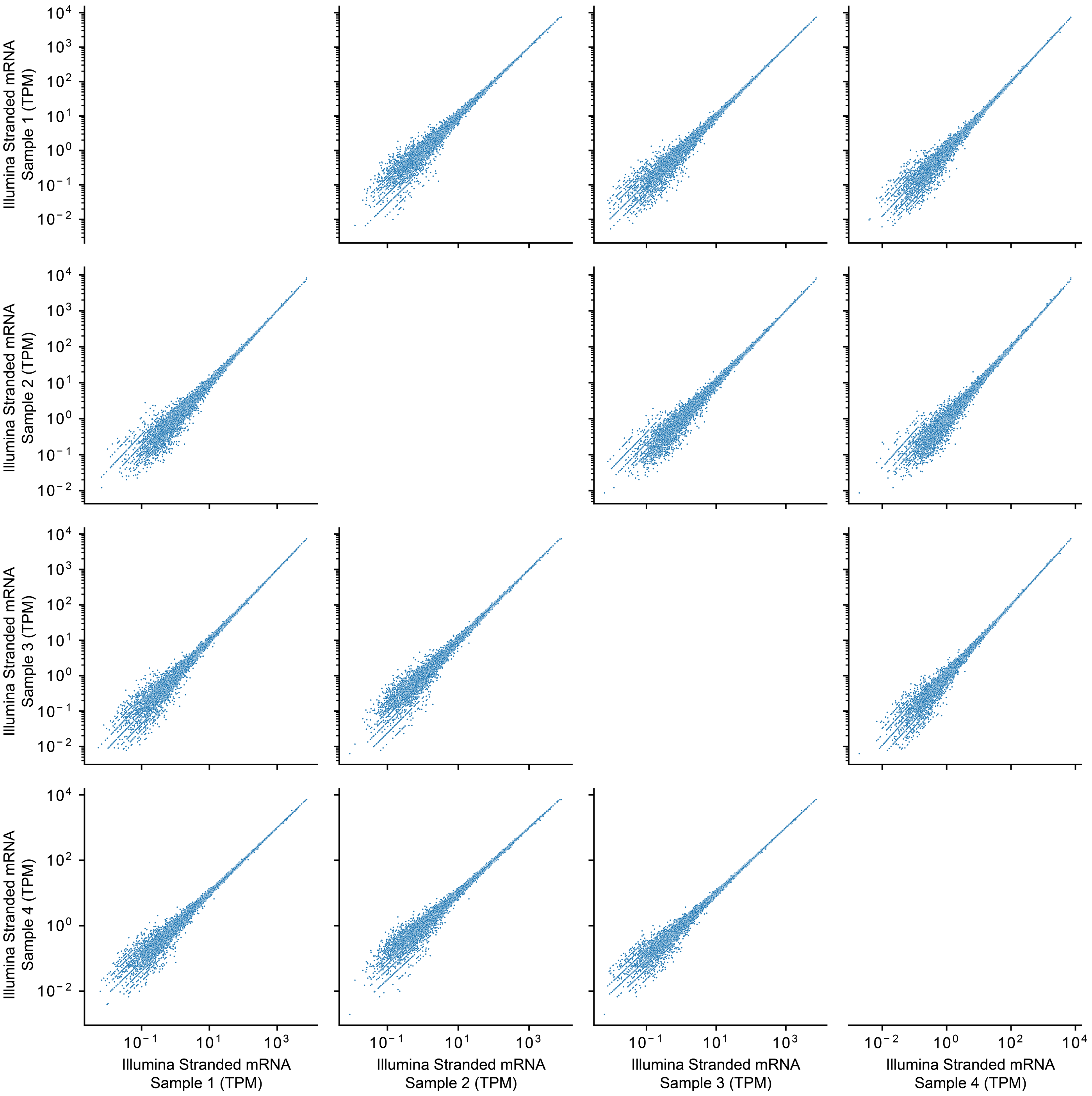


**Figure S4: Illumina Stranded mRNA data shows high intra-method correlation.**
Scatter plots of transcript per million (TPM) counts computed for four Illumina Stranded mRNA experiments (HEK293FT RNA).

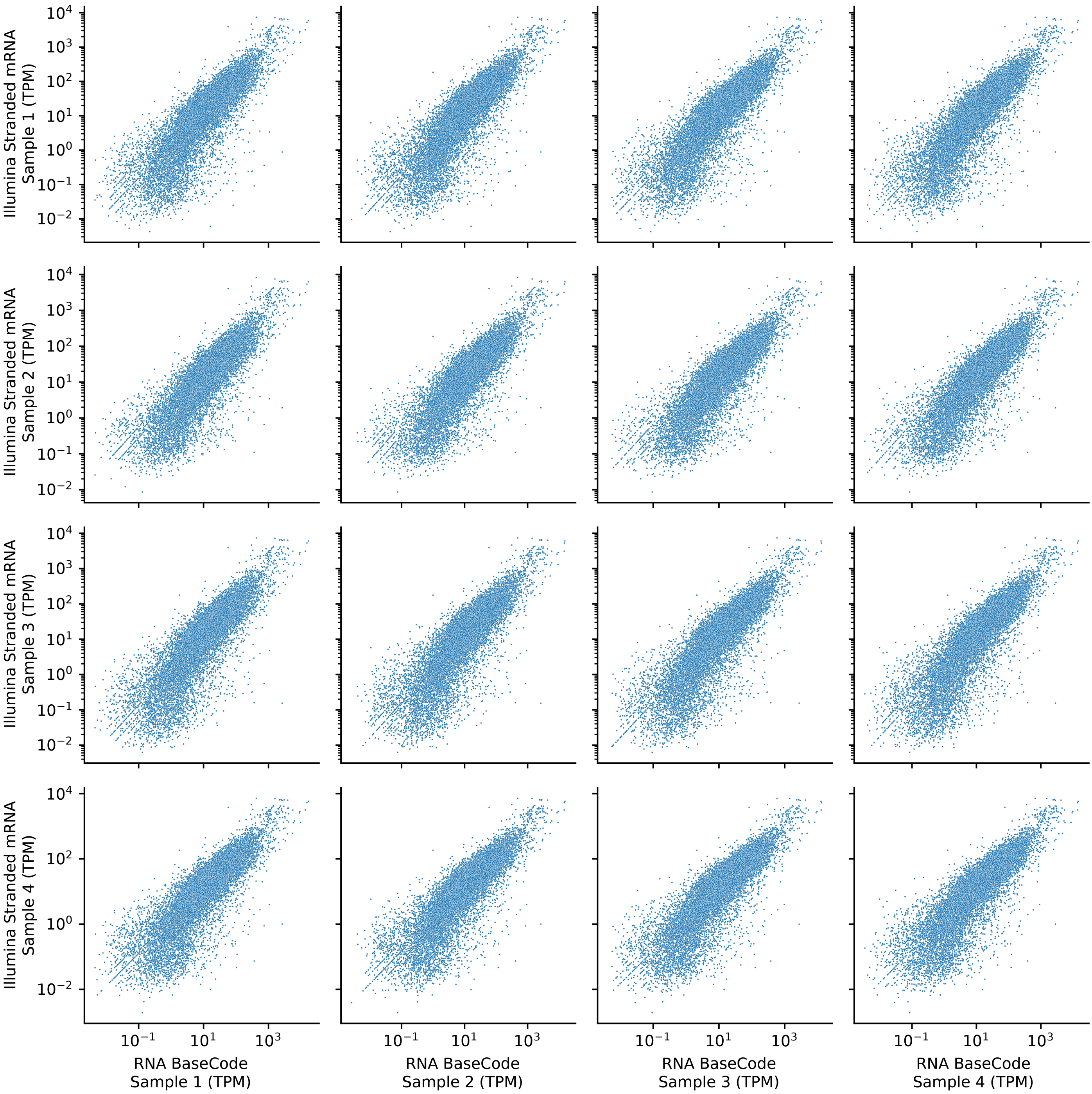


**Figure S5: Scatter plot matrix, comparing RNA BaseCode and Illumina Stranded mRNA data.**
Scatter plots of transcript per million (TPM) counts computed for four RNA BaseCode experiments (x-axis) against TPM counts in four Illumina Stranded mRNA experiments (y-axis).

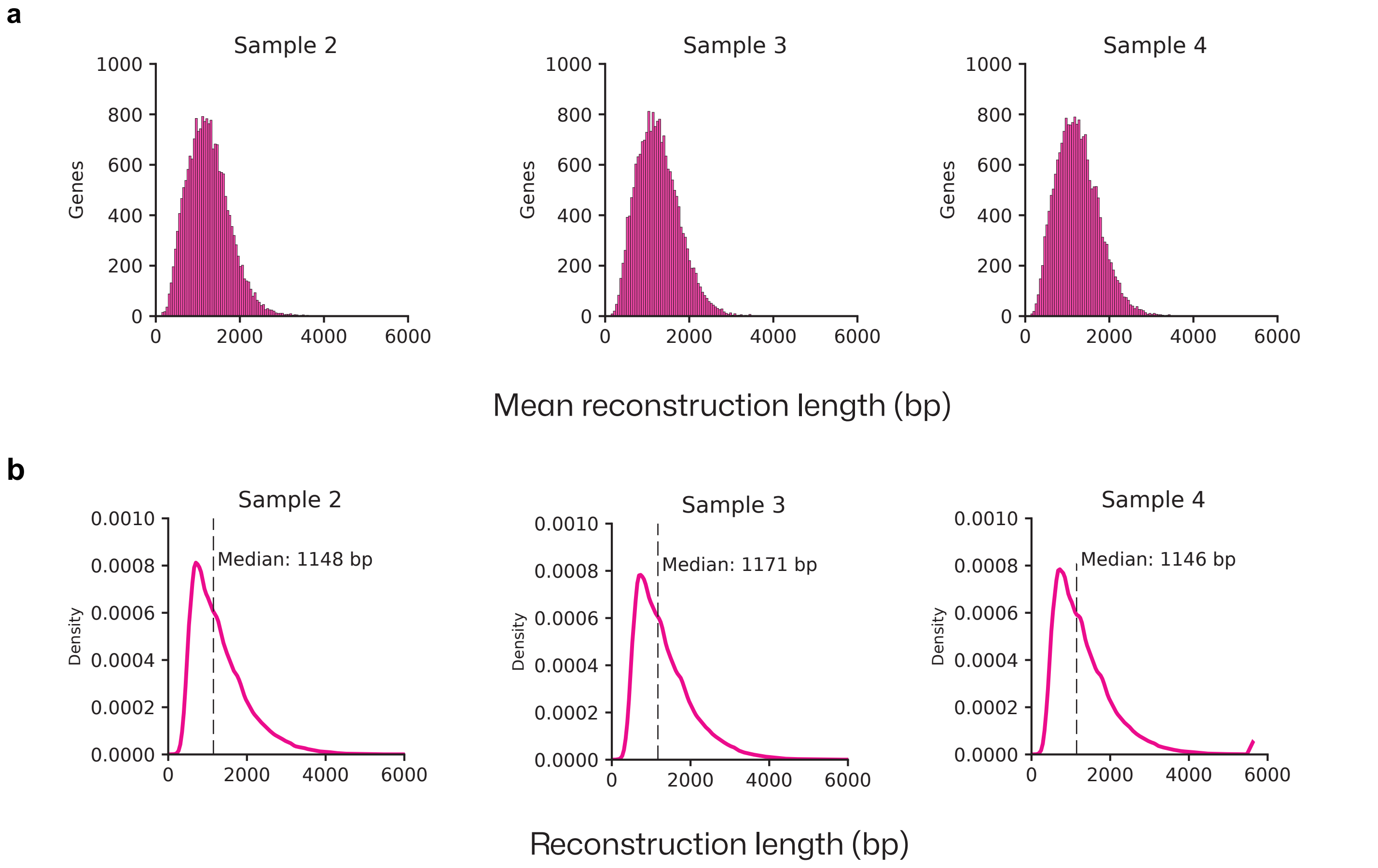


**Figure S6: RNA BaseCode length distribution.**
**(a)** Barplot of the mean length of completed reconstructed molecules across n = 29,494, 29,980 and 29,878 genes for Sample 2, Sample 3 and Sample 4 respectively. For sample 1 see Figure 2h. **(b)** and the length distribution of the individual completed reconstructed molecules for Sample 2, Sample 3 and Sample 4 respectively. For sample 1 see Figure 2i.

a

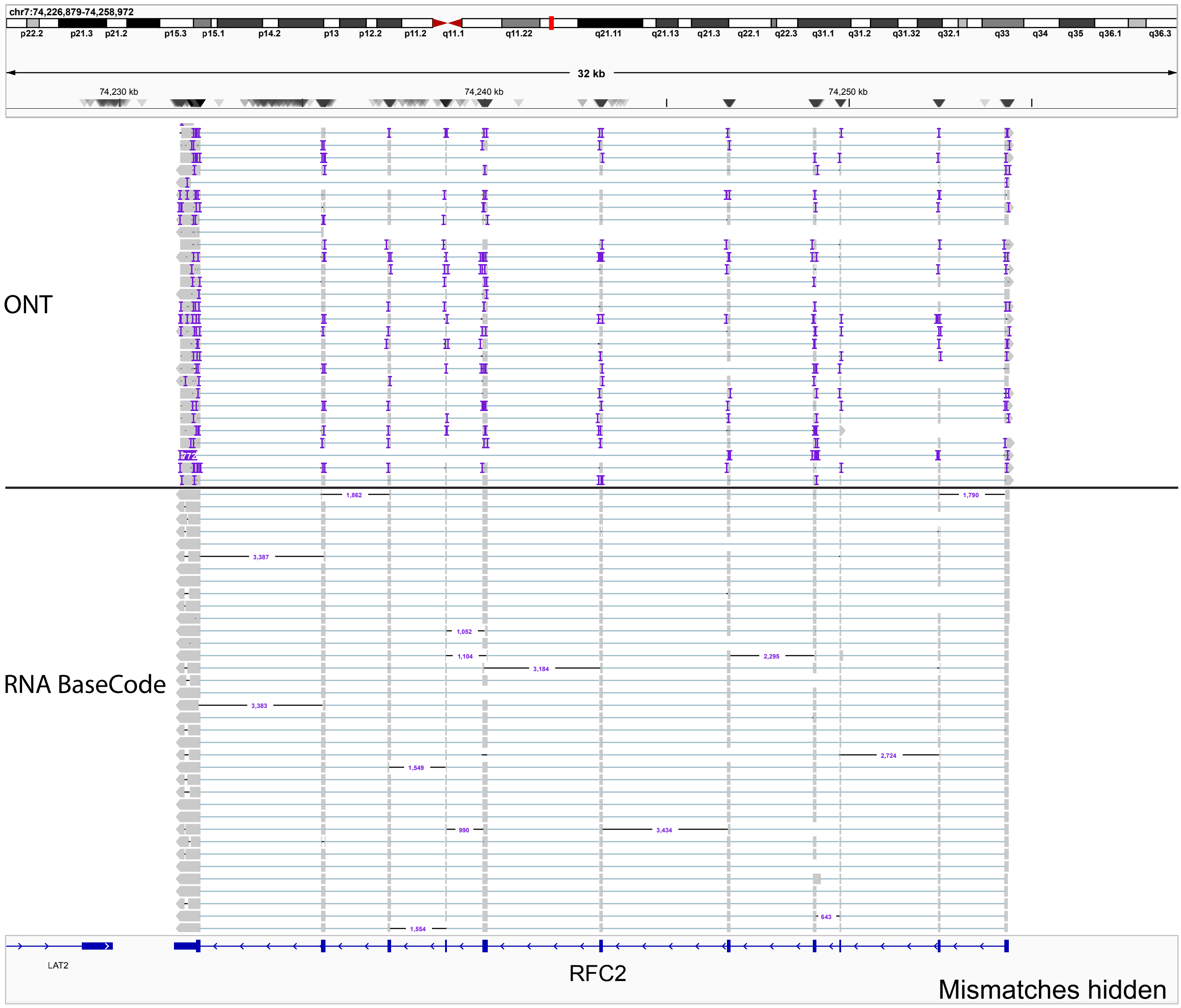


b

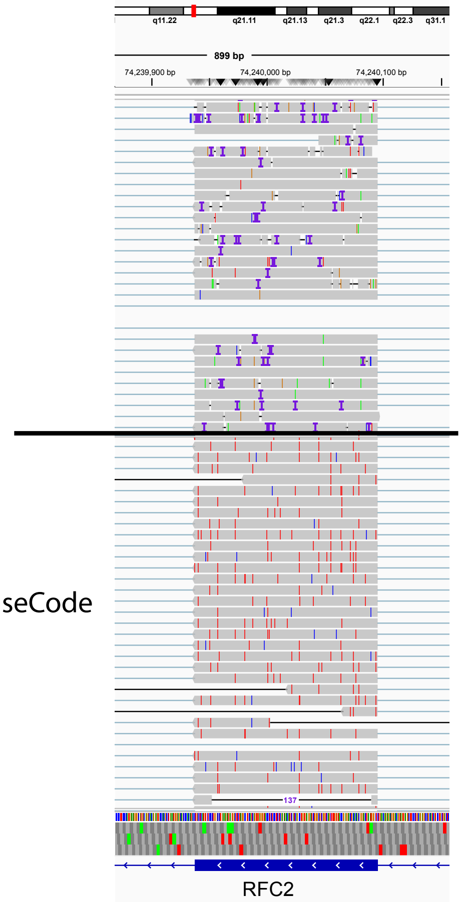


**Figure S7: BaseCode reconstructed sequences are devoid of insertions and deletions (a)**Visualizing small insertions and deletions (indels) present in ONT long-read data (top in each panel) and RNA BaseCode reconstructed data (bottom in each panel) for RNA molecules originating from the RFC2 gene. Mismatches are hidden in both ONT and RNA BaseCode data. The deletions shown for RNA BaseCode indicate missing information, not actual deletions propagated from the short reads. Please see Materials and Methods for more information. **(b)** Same as in S7a, but with mismatches shown in both ONT and RNA BaseCode data. The deletions shown for RNA BaseCode indicate missing information, not actual deletions propagated from the short reads. Please see Materials and Methods for more information.
